# BRED: bioluminescence energy transfer to dye for monitoring ceramide trafficking in cell

**DOI:** 10.1101/2021.03.31.437878

**Authors:** Gita Naseri, Christoph Arenz

## Abstract

Bioluminescence resonance energy transfer (BRET) is a genetically encoded proximity-based tool to study biomolecular interactions. However, conventional BRET is usually restricted to only a few types of interactions like protein-protein or protein-ligand interactions. We here developed a spatially unbiased resonance energy transfer system, so-called BRED - bioluminescence resonance energy transfer to dye. BRED allows transferring energy from a genetically encoded bright human optimized luciferase to a fluorophore-labelled small molecule. The high efficiency of the system allows RET without specific interaction of donor and acceptor. Here, we applied BRED to monitor the trafficking of the signalling lipid ceramide, to the Golgi. This was enabled by an engineered Golgi-resident luciferase, which was used to sense the influx of BODIPY-labeled ceramide into the surrounding membrane. We demonstrated the implementation of the method via flow cytometry, thereby combining the sensitivity of bulk cell methods with the advantages of single-cell analysis. This toolbox enables simple and robust live-cell analysis of inhibitors of CERT-mediated ceramide transport. The design principle of our optogenetic tool can be applied to study intracellular trafficking of metabolites and screen for inhibitors of their key enzymes.

## Introduction

Cellular function is the result of multiple dynamic interactions between different biomolecules. Yet, characterizing these interactions within the living cells is technically challenging [1]. Bioluminescence resonance energy transfer (BRET) [2–5] and fluorescence resonance energy transfer (FRET) [6–8] have been frequently used for this purpose. However, despite numerous examples, these techniques are limited by low brightness and different maturation speed of RET donors and acceptors [9], poor spectral overlap [10], and a low dynamic range of resonance energy transfer (RET) from a donor to an acceptor within a distance of 2−10 nm [11]. These challenges are even more severe when large multicomponent complexes need to be analysed [12] or donor and acceptor distances within a membrane are variable. In the last decade, numerous attempts to overcome the aforementioned challenges have been described, including tailored implementation of organic dyes to increase stability, brightness, and optimized excitation and emission wavelengths of the donor [13, 14] or acceptor [15]. Likewise, novel substrates of the various luciferases have been synthesized, with improved bioavailability, stability, and alternative spectral properties [5, 16–19]. To push the limit of resonance energy transfer measurements beyond 10 nm, optimized zero-mode waveguide aperture structure in combination with molecular constructs featuring multiple acceptor dyes [20] or farFRET allowing enhanced energy transfer using multiple acceptors [12] were recently developed.

Conversely, increasing the number of donors could be another solution to this problem and has been addressed, e.g. by developing codon-optimized luciferases for optimal expression in eukaryotic cells [3, 21, 22]. Another way to increase BRET donor expression may be realized by the use of strong promoters such as human cytomegalovirus (*CMV*) promoter [23]. Up to now, most of the characterized BRET systems have been established on episomal plasmids [2, 3]. A consequence and a major caveat to available BRET systems is the clonal variation of plasmid-based systems, causing a high degree of phenotypic heterogeneity that limits achieving high-level expression, prediction of values, and process streamlining [24]. In contrast, chromosomal integration of the gene of interest (GOI) typically leads to its higher expression due to less metabolic burden and higher homogeneity [25, 26]. However, approaches for the generation of stable human cell lines are usually based on random integration of the GOI into the genome that also leads to clonal variation [24]. To overcome this obstacle, Shin *et al*. (2020) characterized the adeno-associated virus site 1 (*AAVS1*) locus as a genomic safe harbour (GSH) that shows a high level of transcription and protein expression with high homogeneity in engineered human embryonic kidney (HEK293) cells [25].

Here we present a novel tool bioluminescence resonance energy transfer to dye-labelled metabolites, termed BRED, which was designed to extend the applicability of BRET to various types of biomolecular interaction. We sought to track and quantify cellular trafficking of small molecule metabolites labelled with common low molecular weight dyes like BODIPY FL [27, 28]. Organic dyes are small enough to limit the perturbation of the biological processes and can therefore be better suited as biomolecule markers, compared to fluorescent proteins [29]. Our BRED assay relies on a human optimized *Renilla* luciferase (hRLuc) [3], which catalyses the oxidation of coelenterazine-h (CTZ-h) luciferin, a blue-shifted substrate of hRluc with bright luminescence signal [30, 31], [16, 17], to yield relatively stable light emission peaking at 480 nm able to excite a physically unlinked acceptor.

As a proof of concept, we applied BRED for visualizing the transport of the low molecular weight lipid ceramide from the endoplasmic reticulum (ER) to the Golgi [32]. The non-vesicular transport of this lipid is mediated by the ceramide transfer protein (CERT) [33] and is rate-limiting for the biosynthesis of the major plasma membrane component, sphingomyelin [34]. Specific inhibitors of CERT-mediated ceramide transport [35] are considered promising for combating cancer-associated multi-drug resistance [36], as well as different infectious diseases [37, 38]. In our recent study, several potential inhibitors of ceramide intracellular transport were identified using the fluorescence-labelled ceramide (BODIPY-Cer) in a liposomal FRET assay[39]. However, most developed CERT inhibitors have not been tested in cells[40], mainly due to the lack of suitable assays. Inhibitors are usually validated in cells indirectly by inhibition of sphingomyelin biosynthesis using quantitative lipid mass spectrometry [39]. Alternatively, ceramide transport can be evaluated by confocal microscopy, but even super-resolution microscopy is often insufficient for a clear differentiation of Golgi from ER membranes [41]. Noteworthy, setting up such a system would require highly efficient RET between molecules that do not interact with each other directly and whose mean distance is therefore much larger than in classical ligand-receptor interactions. We, therefore, reasoned that monitoring CERT-mediated ceramide transport would be a formidable task to highlight the power of our BRED strategy. Towards this end, a luciferase fused to the CERT-derived Golgi-targeting pleckstrin homology (PH) domain [42] was stably integrated into the *AAVS1* locus of HEK293 cells [25]. We showed the fusion hRLuc enables RET to BODIPY FL-ceramide (BODIPY-Cer), a well-known Golgi stain [32], in either a plate reader format or by using flow cytometry that we adopted for RET study. We further validated that successful RET from hRLuc to BODIPY-Cer in presence of ceramide trafficking inhibitor HPA-12[35]. The concept of BRED tool, together with the experimental evaluation approach, can be applied to evaluate potent drug-like inhibitors of the key enzymes of intracellular transport pathways to speed-up drug development.

## Materials and methods

### General

Plasmids were constructed by NEBuilder HiFi DNA assembly (New England Biolabs, Frankfurt am Main, Germany) or by ligation using T4 DNA (New England Biolabs, Frankfurt am Main, Germany). Plasmid and primer sequences are given in Tables S1 and S2, respectively. PCR amplification of DNA fragments was done using high-fidelity polymerases: Phusion Polymerase (Thermo Fisher Scientific), Q5 DNA Polymerase (New England Biolabs, Frankfurt am Main, Germany), or PrimeSTAR GXL DNA Polymerase (Takara Bio, Saint-Germain-en-Laye, France) according to the manufacturer’s recommendations. All restriction enzymes were purchased from New England Biolabs (Frankfurt am Main, Germany). Amplified DNA parts were gel-purified prior to further use. The primers were ordered from Biomers.net (Ulm, Germany). Plasmids were transformed into *Escherichia coli* ER2925, NEB 5α, or NEB 10β cells (New England Biolabs). Strains were grown in Luria-Bertani medium with an appropriate selection marker at 37°C (Ampicillin, 100 μg mL^−1^ and Kanamycin, 50 μg mL^−1^). The plasmid constructs were confirmed by sequencing (Microsynth Seqlab, Goettingen, Germany).

### Construction of plasmids

pGNPH: The DNA coding sequence (CDS) of PH domain [42] was amplified by PCR (primers GNCA001 /GNCA002, on pHis6-GB1 harbouring CERT PH domain). The ~5000-bp of *Age*I-HF/*Not*I-HF-digested plasmid RasBRET-Tk [3] was assembled with the PCR fragment to generate plasmid pGNPH.

pGN001: The plasmid pGNPH served as a template to PCR-amplified fragment containing *CMV* promoter, PH fused to hRLuc, SV40 polyA signal (primers GNCA003 /GNCA004). The fragment was cloned into *Hin*cII/*Xba*I-digested plasmid AAVS1-LP [25]. The generated plasmid was named pGN001.

pGN002: The ECFP encoding fragment was amplified by PCR using primers GNCA005 and GNCA006 (on pcDNA3-CFP, Addgene #13030), followed by *Dpn*I treatment. Additionally, primer GNCA005 harbors the C-terminal sequence of a Golgi targeting tag [43] of FLAG-tagged mTOR [44]. To add the N-terminal sequence of the Golgi targeting tag [43] upstream of ECFP, the fragment was amplified by PCR using primers GNCA007 and GNCA006. The resulting fragment was digested with *Age*I and *Bam*HI, followed by *Dpn*I treatment. Subsequently, the digested fragment was assembled into ~4000-bp *Age*I/*Bam*HI-digested plasmid RasBRET-tK [3]. The generated plasmid was named pGN002.

### Cell culture

The cells were cultured in Dulbecco’s modified Eagle’s medium (DMEM, Thermo Fisher Scientific) supplemented with 10% FCS (FCS, Thermo Fisher Scientific), 100 U ml^−1^ penicillin, and 0.1mgml^−1^ streptomycin (Thermo Fisher Scientific) at 37 °C with 5% CO_2_. Appropriate number of the cells were trypsinized prior to perform the experiments.

### Construction of BRED 0.1 cell line

HEK293E cells were cultivated as monolayer cultures in a T-flask (Thermo Fisher Scientific) with a working volume of 5 mL at 37 °C with 5% CO_2_ and passaged every 3 days. Stable cell pools that express PH-hRLuc fusion at *AAVS1* locus constructed by transfecting 2500 ng DNA consists of the PH-hRLuc fusion-donor and sgRNA-Cas9 vector at a ratio of 1:1 (w:w) using Lipofectamine LTX (Thermo Fisher Scientific) in 700 μl Opti-MEM was added per well for 4h. After adding 2 mL of DMEM (10% FCS), the cells were incubated overnight at 37 °C with 5% CO_2_. followed by 2 weeks of selection with 3 μg mL^−1^ of puromycin (Thermo Fisher Scientific). Then, cell clones were generated by plating cell pools at 0.3 cells per well. The genomic DNA was extracted using PureLink Genomic DNA Mini Kit (Thermo Fisher Scientific) and confirmed by PCR. The viable cell concentration (VCC) was estimated using a CountessII FL automated cell counter (Thermo Fisher Scientific) and the trypan blue dye exclusion method. Long-term culture for assessing the stability of transgene was done for approximately 1 month. The bioluminescence output at passage 0 and 30 was measured by flow cytometry. Cells were passaged every 3 days in a 6 well plate containing 3 mL culture medium. At every passage, the initial VCC was 2 × 10^5^ cells per ml.

### Plate reader analysis

The bioluminescence measurement was performed for the cell lines HEK293, HEK293 with pGNPH, and BRED 0.1. For HEK293 cells with pGNPH, the cells at 70% confluence were transiently transfected using Lipofectamine 2000 (Thermo Fisher Scientific) and plated on 0.01% poly-d-lysine white 96-well plates at a density of 1×10^5^ cells per well. For transfection, 100 ng vector and 1 μl Lipofectamine 2000 (Thermo Fisher Scientific) in 200 μl Opti-MEM were added per well for 4h. After adding 100μl of DMEM (10% FCS), the cells were incubated overnight at 37 °C with 5% CO_2_. The bioluminescence measurements were performed after 24h of transfection on white 96-well plates. Before measurements, the cells were switched to serum-free medium (DMEM with 0.1% BSA) for 5h. Cells were washed with ice-cold phosphate-buffered saline (PBS) and a modified Krebs-Ringer buffer[3] was added to the cell. Additionally, an appropriate concentration of CTZ-h was added to the wells and the cells were incubated for 5min at 37 °C with 5% CO_2_ in darkness. The measurements were performed at 37 °C using plate reader VICTOR X5 (PerkinElmer). The detection time was 0.5 s. The luminescence records were the average of at least three independent experiments. The BRET assay was performed for the cell lines HEK293, BRED 0.1, and HEK293 with pGNPH. The transfection of HEK293 cells with pGNPH was performed as described above. The cells were trypsinized and 1×10^5^ cells were seeded per well for BRET measurements. The BRET measurements were performed after 24h of transfection on white 96-well plates. Before measurements, the cells were switched to a serum-free medium for 5h. Appropriate concentration of BODIPY FL-Cer and /or inhibitor (see **Results and discussion**) were added to the wells and the cells were incubated for 45min at 37 °C with 5% CO_2_ in darkness. Cells were washed with ice-cold PBS and a modified Krebs-Ringer buffer[3] was added to the cell. An appropriate concentration of CTZ-h (see **Results and discussion**) was added to the wells and the cells were incubated for 5min at 37 °C with 5% CO_2_ in darkness. The measurements were performed at 37 °C using plate reader POLARstar OPTIMA (BMG LABTECH), which allows for the detection of signals using filters at 485-and 530-nm wavelengths. The detection time was ~0.45 s. The BRET ratios were calculated as the 530 nm/485 nm ratio. The data records are the average of at least three independent experiments.

### Flow cytometry analysis

BD FACSMelody flow cytometer (BD Biosciences) was used for flow cytometry analysis. 1 × 10^5^ cells were seeded per well of a 6-well. To assess the fluorescent output of BODIPY FL-Cer dye, an appropriate concentration of dye was added (see **Results and discussion**) to the wells after 24h and the cells were incubated for 45min at 37 °C with 5% CO2 in darkness. After trypsinization, cells were washed with ice-cold PBS and a modified Krebs-Ringer buffer was added to the cell. The MFI of the BODIPY FL-Cer was detected using a BD FACSMelody flow cytometer (blue laser, 527/32 nm filter). To avoid measurement error, cell samples were fixed using 70% ethanol, kept at −20 °C, and analyzed. To setup the BD FACSMelody flow cytometer for BRED 0.1 system, an optical configuration for the combination of blue and YG channels was created. 1 × 10^5^ of BODIPY FL-Cer treated HEK293 cells were loaded into the system. Subsequently, the 527/32 nm filter is placed in a channel for yellow-green laser. The fluorescence values were obtained in both channels. To measure the MFI of BRED 0.1 cells treated with BODIPY FL- Cer and CTZ-h, cell suspension in modified Krebs-Ringer buffer was treated with CTZ-h and incubated for another 5min at 37 °C with 5% CO_2_ in darkness. Under the new configuration setup, the MFI of BRED 0.1 cells was detected. The fluorescence values were obtained from a minimum of 10,000 cells per sample. To avoid measurement error, cell samples were fixed using 70% ethanol, kept at −20 °C, and analyzed. The mean fluorescence per cell was calculated using FlowJo software.

### Confocal microscopy

To analyze the localization and distribution of the BODIPY FL-Cer, 8 well μ-slides (ibidi, ibiTreat) were coated with 0.01% poly-d-lysine for 10 min. Subsequently, the solution was removed and the slides were dried. The 1 × 10^4^ HEK293 cells were seeded and incubated in DMEM (10% FCS) overnight at 37 °C with 5% CO_2_. The cells were transfected with 100ng vector (see Plate reader analysis) Appropriate concentration of BODIPY FL-Cer was added (see Results) to the wells and the cells were incubated for 45 min at 37 °C with 5% CO_2_ in darkness. After trypsinization, cells were washed with ice-cold PBS and a modified Krebs-Ringer buffer[3] was added to the cell. The cells were imaged in two different channels (CFP: λ_ex_ = 438/24 nm, λ_em_ 483/24 nm, YFP: λ_ex_=500/24nm, λ_em_: 545/40) using an IX83 microscope from Olympus and the images were processed by OlyVIA.

## Results and discussion

### The workflow of BRED and its application

In the BRED system, the energy donor and acceptor are spatially unbiased. The adequate amount of a hRLuc [3] as the energy donor is positioned within the desired compartment of the cell. This is achieved by generating a donor cassette consisting of hRLuc fused to the organelle targeting domain under the control of a strong *CMV* promoter. The cassette is stably integrated into a GSH locus of the genome, enabling high protein expression with high homogeneity. Upon adding CTZ-h (a bright substrate of hRLuc) [45], the interaction of the donor with a physically-unlinked molecule of interest conjugated to suitable acceptor fluorophore [28], provided in close proximity to the hRluc, can be studied using the flow cytometry protocol introduced in this study (see **Flow cytometry-based monitoring of ceramide trafficking in BRED system**). In view of the crucial role of ceramide for cell fate and cancer development [46], we applied the BRED system to study ceramide trafficking. A donor cassette containing a pleckstrin homology (PH) domain [42] fused to a C-terminal hRLuc was integrated into the *AAVS1* genomic site of HEK293 cells to generate the BRED 0.1 cell line. Green BODIPY-conjugated ceramide (BODIPY FL-Cer) [32] was used as an acceptor. Flow cytometry enabled us to evaluate the transport of BODIPY ceramide to Golgi membranes inside the living cell (Fig. 1).

**Figure 1.**
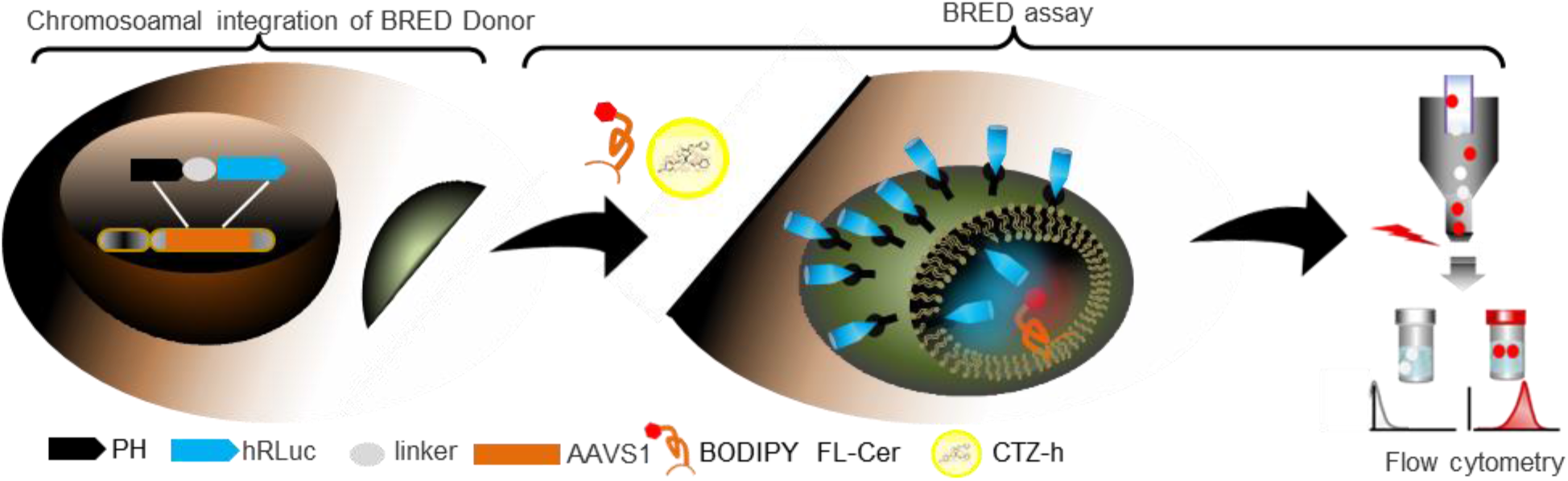
Schematic diagram of BRED to study ceramide trafficking. Chromosomal integration of BRED Donor. A donor cassette expressing the hRLuc linked to the PH domain is integrated into the *AAVS1* locus of the HEK293 genome. BRED assay: Upon adding CTZ-h (hRLuc substrate) and green BODIPY- Cer dye (ceramide acceptor), the resonance energy is transferred from Golgi-anchored Rluc-CTZ donor couple to ceramide. Flow cytometry is used to study ceramide transport across the Golgi membrane of the living cell. PH, pleckstrin homology domain; *AAVS1*, adeno-associated virus site 1; BODIPY FL-Cer, borondipyrromethene fluorescein-ceramide; CTZ-h, Coelenterazine-h; hRLuc, human optimized *Renilla* luciferase.

### Characterizing BRED donor and acceptor for ceramide trafficking

To generate the genetically encoded BRED donor, the plasmid pGNPH (Fig. 2a) encoding hRLuc [47] fused to the N-terminal PH domain of CERT [42] through an 18-aa spacer [42] was generated for transient expression. For stable expression, the BRED donor was integrated into the *AAVS1* locus of HEK293 cell using clustered regularly interspaced short palindromic repeats (CRISPR)/CRISPR-associated protein (Cas9) [25] and single-guide RNA (sgRNA) targeting the intron between exon 1 and exon 2 [48](Fig. 2b). The generated cell line was named BRED 0.1. Bioluminescence intensity in BRED 0.1 cells (black line) or HEK293 cells transfected with pGNPH (pink line) treated with 10 μM of CTZ-h, or in BRED 0.1 cells treated with 5 μM of CTZ-h (red line) was evaluated (Fig. 2c). HEK293 cells without substrate treatment (grey line) were used to observe the cell-alone signal. In the case of BRED 0.1 cells treated with either 5 μM or 10 μM of CTZ-h, a significant bioluminescence intensity was observed that gradually decreased with time and after 7 minutes was similar to that of HEK293 control cells. In contrast, the bioluminescence intensity observed for HEK293 cells transfected with pGNPH was slightly (but not significantly) higher than that of HEK293 cells, only during the first 3 minutes of time-course measurement. The RLuc-CTZ-h donor couple leads to a maximum emission at 480 nm[49], a wavelength that is appropriate for excitation of BODIPY FL, which subsequently reemits light at 530 nm (Fig. 2d). When administered into the cell medium, BODIPY FL ceramide (BODIPY-Cer) was able to stain unmodified HEK293 cells efficiently, as evaluated by flow cytometry (Fig. 2e, CV=89.1). Next, cells were transfected with pGN002 (see **Material and methods**), for Golgi expression of enhanced cyan fluorescent protein (ECFP), and Golgi localization of the BODIPY-Cer in HEK293 cells was confirmed by confocal microscopy (Fig. 2f).

**Figure 2.**
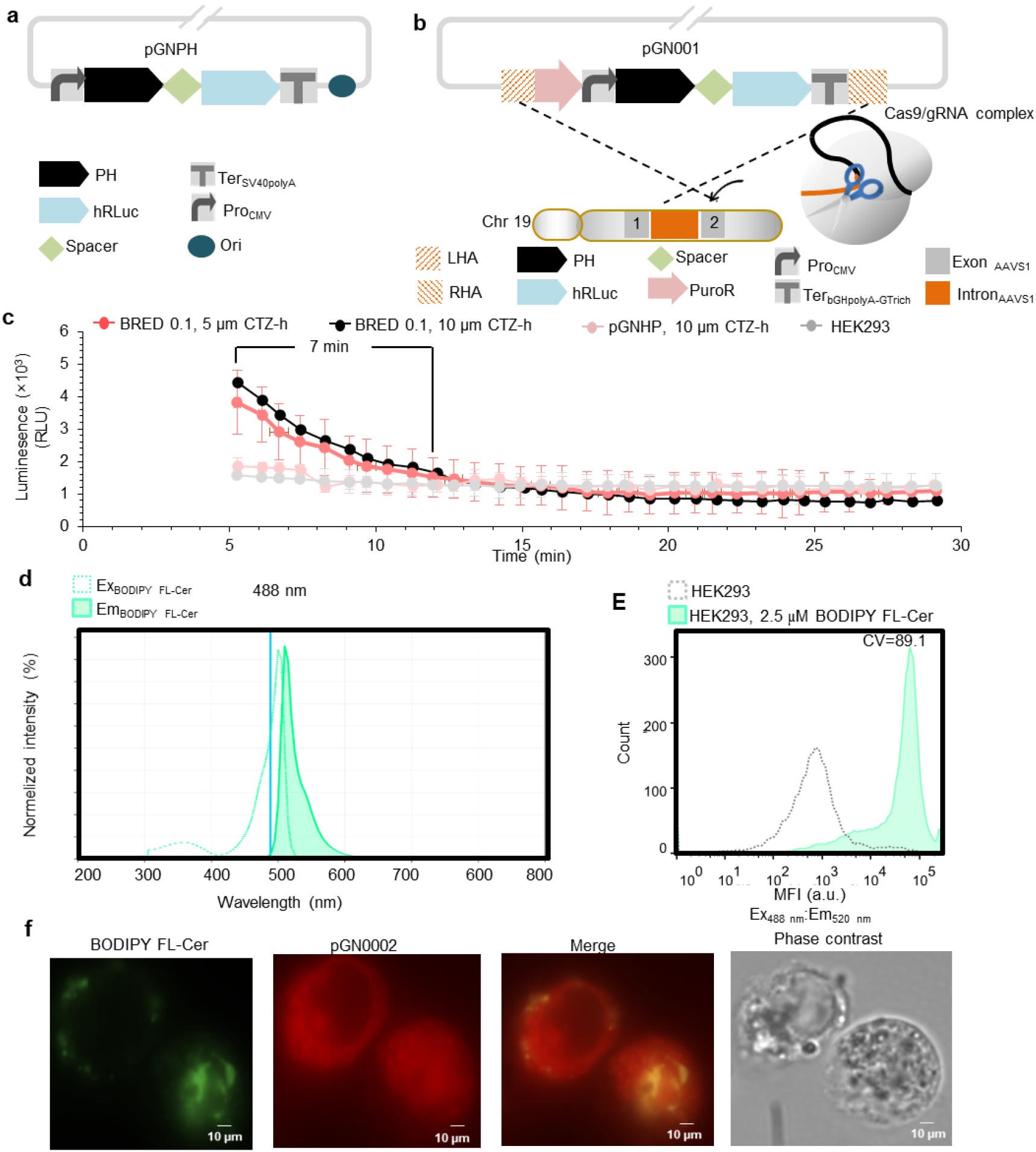
Evaluation of BRED donor and acceptor partners for ceramide transport. a, A schematic representation of the generation of stable HEK293 cell lines harbouring energy donor. The donor plasmid pGN001 contains *CMV* promoter, *hRluc*, 18-bp spacer, *PH*, bGH poly*(A)-GU*-rich terminator, flanked by the 900 bp homology arm left and right arms. A DNA double-strand break induced by sgRNA and Cas9 protein was repaired by homology-directed repair using the donor fragment that targets the *AAVS1* locus located on chromosome 19 [25]. **b**, A schematic representation of the plasmid pGNPH. The plasmid contains *CMV* promoter, *hRluc*, 18-bp spacer, *PH*, and *SV40* poly(A) termination signal. **c**, Time course measurement of the luminescence intensity in the living cell. BRED 0.1 cells (black line) were exposed to 10 μM CTZ-h, while HEK293 cell (grey line), HEK293 cell with episomal plasmid pGNHP (pink line), and BRED 0.1 cell (red line) were exposed to CTZ-h substrate at 5 μM. After five minutes substrate incubation, the luminescence was measured at intervals of approximately 45 s for 25 min. RLU, relative light unit. Data are means ± SD from three biological replicates. **d**, Spectral characteristics of BODIPY-FL dye. Normalized fluorescence emission (dashed cyan line) and excitation spectra (solid cyan line) of BODIPY FL dye were shown. The maximum emission spectrum of the hRLuc-CTZ-h donor couple (blue line) [31] has overlap with the excitation spectrum of the acceptor. e, Flow cytometric histogram overlay of a BODIPY FL-Cer treated cells onto the control cells. HEK293 cells treated with 5 μM BODIPY-Cer (cyan curve) were analysed in the green channel (Em_488 nm_: Ex_527 nm_) of FACS. CV value of 89.1 was obtained by flow cytometry. The HEK293 cells without treatment were used as a control population (dashed grey curve). FACS histogram is representative data. The fluorescence intensity and CV value obtained from three independent cultures and determined in three technical independent experiments. MFI, mean fluorescent intensity; CV, coefficient of variation; a.u. arbitrary units. **f**, Monitoring the transfer of BODIPY-Cer to the Golgi. Confocal images show the distribution of BODIPY FL-Cer (green) and Golgi apparatus costaining (pGN002; red) in HEK293 cells. Figures show representative images. The experiment was performed in three biological replicates with similar results. Scale bars, 10 μm. BODIPY-Cer, borondipyrromethene FL fluorescein-ceramide; CMV, cytomegalovirus; CTZ-h, Coelenterazine-h; hRLuc, human optimized *Renilla* luciferase. Full data for Fig. 2c, and 2e are given in Data S1a and S1b, respectively.

### RET performance of BRED for ceramide trafficking

After characterization of the donor and acceptor partners of our BRED strategy (Fig. 2), the RET performance of BRED (ratio of emission intensities at 520 nm and 480 nm) was evaluated for different cell lines after adding the donor substrate in the presence or absence of BODIPY-Cer acceptor (Fig. 3). In BRED 0.1 cells (the *AAVS1*-integrated hRLuc-PH fusion donor) devoid of BODIPY-Cer, the addition of the CTZ-h substrate led to a signal emission at 480 nm, but not at 520 nm, and thus an acceptor/donor intensity ratio less than 1 was observed (Fig. 3, blue line), as expected (see Fig. 2c, red line). This confirmed the selectivity of the optical windows for CTZ-h (480 nm) and BODIPY (520 nm) emission, respectively. We further showed that treating HEK293 cells with either CTZ-h or BODIPY-Cer does not affect the signal emission at 480 nm and 520 nm (Supplementary Results and Fig. S1). In BRED 0.1 cells, in presence of BODIPY-Cer, a strong energy transfer and thus a 480 nm/520 nm emission ratio > 1 was observed within 5 min after adding CTZ-h substrate. By increasing the concentration of acceptor from 2.5 μM (Fig. 3, orange line) to 5 μM (Fig. 3, red line), an even higher RET signal was obtained for each acquisition time between minutes 5 and 10, with netBRET values (intensity ratio of emission at 520 nm and 480 nm minus the same ratio in absence of acceptor) [50] of ~2 at minute 5. These values dramatically decreased to the level of noise signal of BRED 0.1 cells (Fig. 3, grey line) and HEK293 cells (Fig. 3, pink line). A high donor to acceptor ratio may generally lead to a non-specific RET signal, due to incomplete saturation of donors with acceptors [51]. We observed a non-linear change of the RET signal while increasing the amount of the acceptor, suggesting better saturation of all donors with acceptor molecules at higher concentrations [51]. Taken together, our result suggested that chromosomally expressed PH-hRluc fusion in presence of 5 μM of CTZ-h substrate enables efficient RET to BODIPY-Cer, which is in the same subcellular compartment.

**Figure 3.**
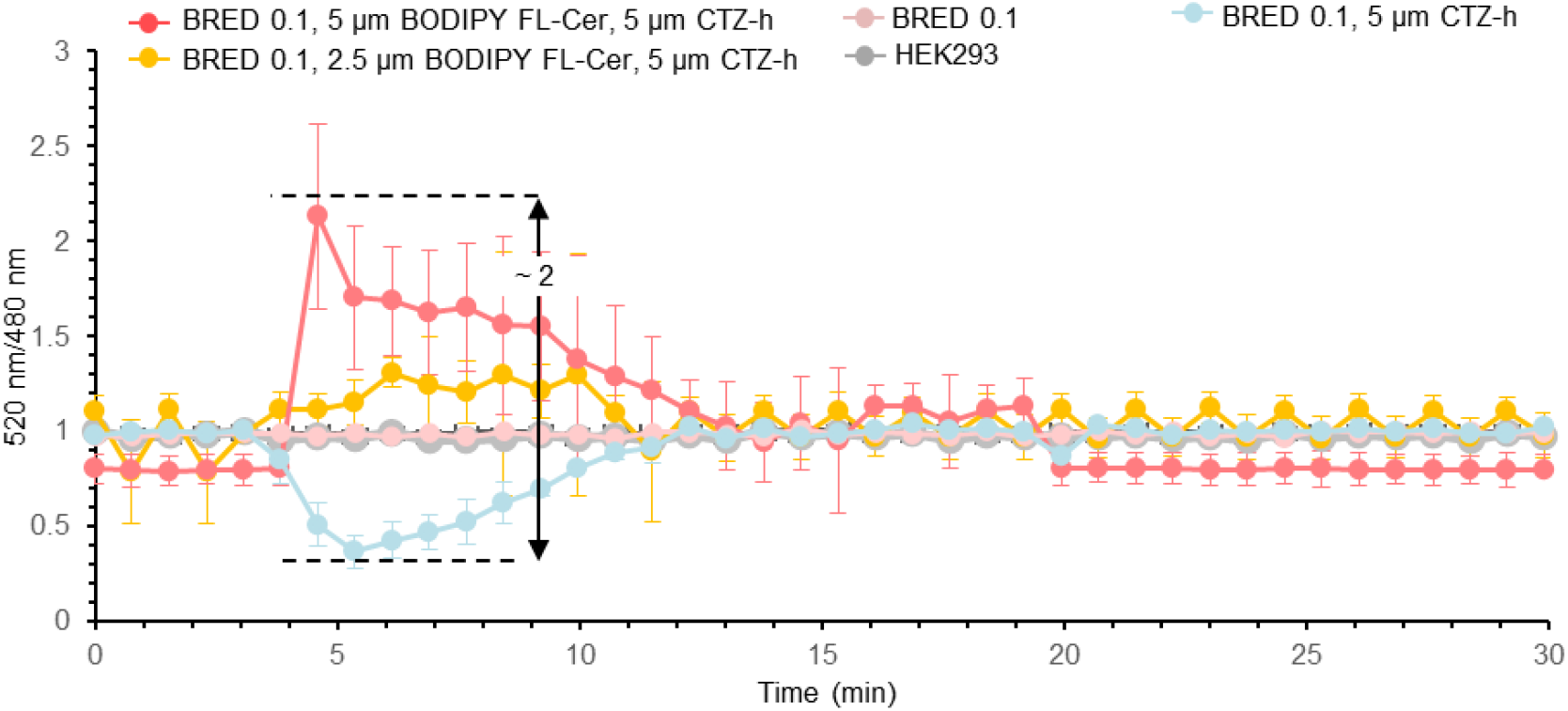
Resonance energy transfer of BRED for ceramide transport over time. Average 535nm/480nm ratio intensity measured every 45 s for 30 min for BRED 0.1 (pink line) and HEK293 cells without CTZ-h and BODIPY FL-Cer treatment (grey line), the BRED 0.1 cells treated with only 5 μM of CTZ-h (blue line), BRED 0.1 cells treated with 5 μM of CTZ-h and BODIPY FL-Cer at 2.5 μM (orange line) or 5 μM (red line, netBRET[50] of ~2). Data are means ± SD from three biological replicates. BODIPY-Cer, borondipyrromethene FL fluorescein-ceramide; CMV, cytomegalovirus; CTZ-h, Coelenterazine-h; hRLuc, human optimized *Renilla* luciferase. Full data are given in Data S2.

### Flow cytometry-based monitoring of ceramide trafficking in BRED system

The RET is often studied using the microscope, which means it can be difficult analysing large numbers of cells in one setting. In addition, luminescence-based RETs generate too dim signals, hard to detect using conventional microscopes. A conventional widefield microscope was successfully applied for bioluminescence imaging of a highly bright Nanoluc [50]. However, the luminescence emitted by Nanoluc is not a well-suited donor for BRED system. Using standard widefield imaging equipment, we showed that hRluc-CTZ-h donor couple could be used to track proteins of interest in BRED 0.1 cells (see Supplementary Methods for the protocol, Supplementary Results, Fig. S2a). Moreover, BRED 0.1 cells treated with CTZ-h and the acceptor showed a higher signal, compared to BRED 0.1 cells exposed to only CTZ-h (Supplementary Results and Fig. S2b). However, the signal was weak, possibly due to the general limitation of the conventional widefield microscope to detect BRED bioluminescence signals.

Therefore, we sought to utilize flow cytometry to evaluate the BRED system. In an earlier report, flow cytometry analysis was implemented to characterize bioluminescence signals emitted by various luciferases-substrates donor couples in insect cells [52]. However, mainly due to the short half-life of the tested donor couples (maximum ≈ 145 s for click beetle luciferase), only a small percentage of cells (16.8% for click beetle luciferase) produced the luminescence signal [52]. In a flow cytometry setup, we omitted the undesired emission fluorescence signal of the BODIPY dye (at 527 nm) due to the excitation at 488 nm in the channel for blue laser by placing the 527/32 nm filter in the channel for yellow-green (YG) laser (Supplementary Results and Fig. S3a and 3b).

This setup allowed evaluating the RET from luminescence from highly expressed hRLuc to BODIPY-Cer to the Golgi in the absence and presence of ceramide inhibitor. Using our protocol, we evaluated the BRED output by flow cytometry (Fig. 4). The overlay histograms presented in the left panel of Fig. 4 illustrate the fluorescent profiles at 527 nm (BODIPY emission), and the graphs presented in the right panel of Fig.4 display the average mean intensities (at 527 nm) obtained from nine replicates of various treatments. BRED 0.1 treated with CTZ-h and BODIPY-Cer (Fig. 4a, green) showed ~1.4-fold higher mean fluorescent output than the control HEK293 cells (Fig. 4a, dashed grey). In contrast, no significant increase in mean intensity was observed for the cells expressing hRluc alone (Fig. 4a, blue), or in combination with either CTZ-h substrate (Fig. 4a, orange) or BODIPY-Cer acceptor (Fig. 4a, red) over the control untreated HEK293 cells (Fig. 4a, dashed grey). The sample with high (Fig. 4a, green) and low (Fig. 4a, dashed grey) fluorescent outputs were used as positive and negative controls, respectively, to analyse the data obtained upon further optimizations. To improve the signal observed for BRED system, we increased the concentration of the acceptor from 2.5 μM to 5 μM. As shown in Fig. 4b (dark green) ~2.5-fold increase of fluorescent output was observed, indicating a dynamic fluorescence output for different local concentrations of BODIPY-Cer. Moreover, we quantified the BODIPY-Cer output derived from transient plasmid-based luminescence expression in presence of a 5 μM acceptor (Fig. 4b, left panel, pink). Although we observed a fluorescent output confirming the resonance energy is transferred from the donor to the acceptor, the majority of cells (81.3%) were exhibited a background signal with the plasmid setup, indicating a high level of heterogeneity among the population (Fig. 4c). Notably, the two populations likely represent cells provided with CTZ-h and BODPY ceramide in the absence (low fluorescence) and presence (high fluorescence) of the hRLuc. To further evaluate the emission observed at 528 nm is due to energy transfer (not re-absorption of the emitted luminescence by the BODIPY dye), and to validate the sensitivity of flow cytometry-based assay, we monitored the effect of ceramide trafficking inhibitor HPA-12 [35, 53] on BRED 0.1 cells. Exposing 5μM of HPA-12 to the BRED 0.1 cells treated with CTZ-h and 5 μM of BODIPY-Cer, the fluorescent output decreased to the level of the donor-alone signal (~4.5-fold less than that of BRED 0.1 cells treated with CTZ-h and 5 μM of BODIPY-Cer).

**Figure 4.**
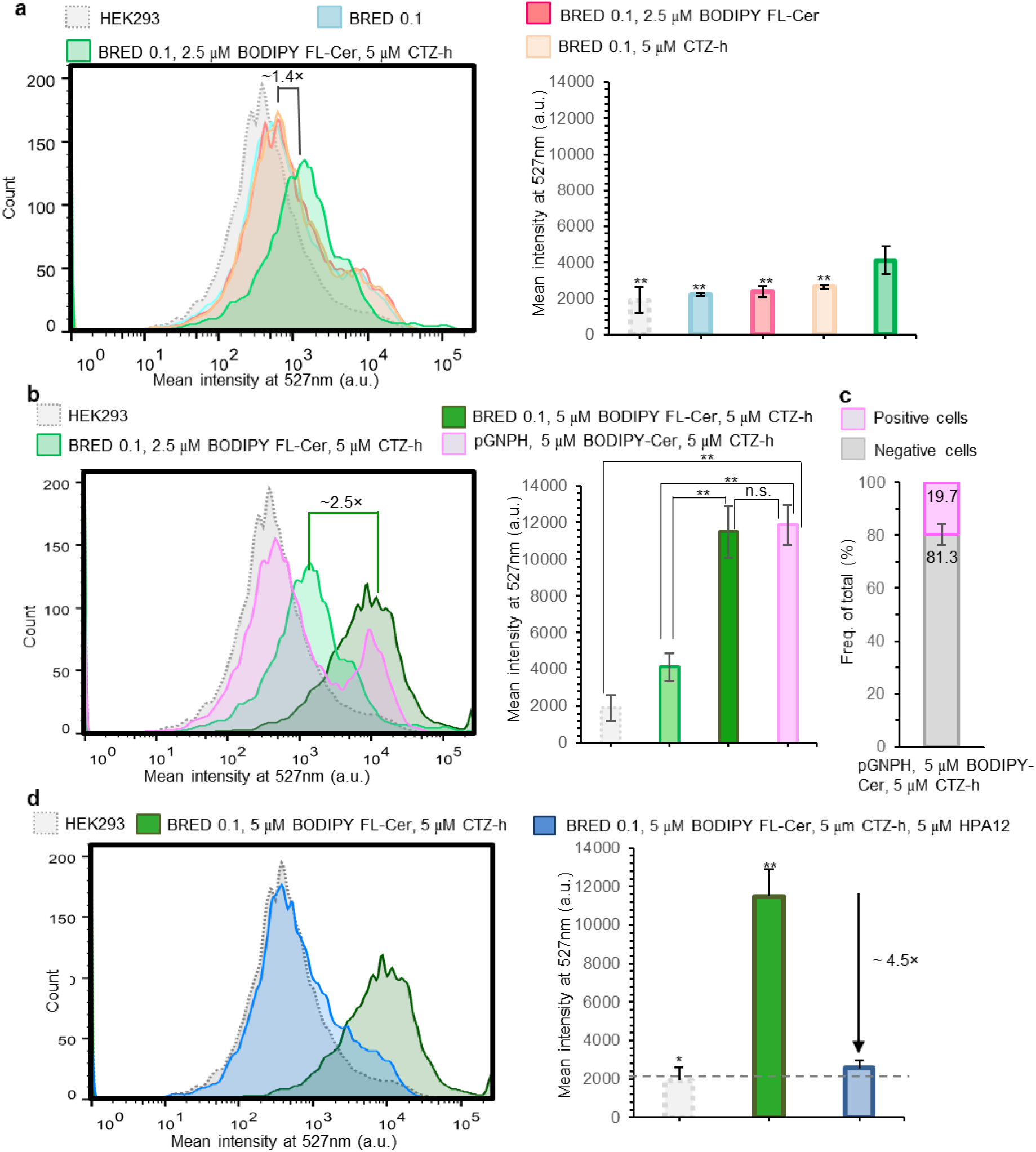
Evaluation of BRED using flow cytometry. In the overlayed histograms, the mean intensity and cell number are indicated on the x- and y-axes, respectively. In the column charts, FACS data are shown that are representative of the mean ± SD of the fluorescence intensity obtained from three independent cultures and determined in three technical independents. a, BRED efficiency analysis. The fluorescent intensity at 527 nm in BRED 0.1 cells stably expressing Golgi-targeted hRLuc in combination with 5 μM of CTZ-h substrate and 2.5 μM of BODIPY FL-Cer showed ~1.4-fold higher fluorescent output (green) than the control HEK293 cells (dashed grey). The cells expressing donor alone (blue), and in combination with the substrate (orange) or acceptor (red) exhibited background level signal. Asterisks indicate a statistically significant difference from hRLuc-CTZ-h pair couple in presence of 2.5 μM of BODIPY FL-Cer (Student’s *t*-test; (∗) *p* < 0.05; (∗∗) *p* < 0.01). **b**, Effect of the acceptor concentration on BRED efficiency. BRED 0.1 (dark green) and HEK293 cells transiently expressing the energy donor (pink) showed 2.5-fold improvement in fluorescent output in presence of 5 μM of CTZ-h, when the concentration of the acceptor increased from 2.5 μM (green) to 5 μM. The HEK293 cells without any treatment (dashed grey) served as a negative control. Asterisks indicate statistically significant difference (Student’s *t*-test; (∗) *p* < 0.05; (∗∗) *p* < 0.01). **c**, Percentage of the positive and negative cell population for plasmid-derived expression of the donor. Only 19.7% of HEK293 cells transfected with plasmid pGNPH (pink) generated the BRED signal and 81.3% exhibited background signal (grey). Freq of the total, Frequency of the population out of the total population. **d**, Effect of HPA-12 [35] on the performance of BRED system. Exposing 5μM of HPA-12 inhibitor to the BRED 0.1 cells treated with 5 μM of CTZ-h and 5 μM of BODIPY FL-Cer (blue) resulted in ~4.5-fold decrease of fluorescent output, compared to BRED 0.1 cells treated with 5 μM of CTZ-h and 5 μM of BODIPY FL-Cer (dark green) and reached to the level of the signal observed for control HEK293 cells (dashed grey). FACS data are representative of the mean ± SD of the fluorescence intensity. Asterisks indicate a statistically significant difference from hRLuc-CTZ-h pair couple in presence of BODIPY FL-Cer and HPA-12 (Student’s *t*-test; (∗) *p* < 0.05; (∗∗) *p* < 0.01). a.u., arbitrary unit, MI, mean intensity; Em_hRLuc_, emission signal of human optimized *Renilla* luciferase; Ex_527 nm_, excitation signal of BRED in 527 nm. Full data for Fig. 4a, 4b, 4c, and 4d are given in Data 2a, 2b, 2c, and 2d, respectively.

### Discussion

The fundamental importance of developing *in vivo* monitoring tools has been widely recognized to understand cell life, which is governed by the dynamics of its molecular actors[54]. The rapid growth of optical techniques based on BRET places them among the most powerful methods to study the dynamics of protein-protein or protein-ligand interactions at the subcellular level [8, 12, 31, 55, 56]. However, our ability to generalize these tools is still in its infancy [50]. It is timely to reconsider the weaknesses of RET systems to encourage promising expansion of their current abilities, as the applications of these methods increase intensely. The improvements may primarily focus on (i) increasing the quantum yield of the donor [57], and (ii) decreasing the artifacts caused by the expression of fusion proteins in BRET systems on protein-protein interactions [31, 51, 58]. Additionally, high-throughput assays to analyse such improved systems would represent a strong asset [52, 59].

The BRED platform presented here has distinct advantages over currently available BRET systems. The exceptionally high efficiency of our system allows efficient energy transfer from a highly expressed luminescence donor to a dye acceptor (with higher photostability and brightness than fluorescent proteins), although a direct interaction between the two has not been described. It was previously reported that clustering of proteins within a membrane leads to increased RET [60]. However, only netBRET values between 0.2 and 0.5 were detected in this study [60], whereas a netBRET value of 2 was obtained for the BRED 0.1 cell line (Fig. 3).

In addition to quantification of RET in our system by conventional bioluminescence assay in a plate reader setup, we show that the BRED system can be linked with flow cytometry. The flow cytometry setup clearly showed the superiority of stable genetic integration over episomal expression and successful application of BRED system for ceramide trafficking study (Fig. 4). Both setups can be used to easily monitor the trafficking of ceramide to the Golgi. The sensitivity for screening ceramide transport inhibitors in living cells is demonstrated using the well-known CERT inhibitor HPA-12 [35], causing a ~4.5-fold decrease in BODIPY fluorescence to virtually background signal (Fig. 4d).

Considering a number of candidate potent drug-like inhibitors that target the ceramide transport to the Golgi [35, 39, 61], the BRED 0.1 cell line would be expected to be convenient for *in cellulo* characterization of their pharmacodynamics. Other organelle targeting domains can be used in lieu of the PH domain to evaluate the transport of ceramide to other cellular organelles, such as mitochondria to study the function of ceramide in regulating mitochondria-mediated cell death [47] and to screen its potent drug-like inhibitors [39, 47] This assay could potentially be expanded to study the intracellular trafficking of other metabolites for which appropriate fluorophore conjugates can be produced. The blue-shifted BODIPLY FL dye in combination with other acceptors such as red-shifted fluorophores can be an advantage when dual BRED monitoring is required to study complex signalling networks [27, 62]. To further optimize BRED system, orthogonal strong synthetic transcription factors [63, 64] can be employed to express the donor protein at the desired time. We believe the BRED reported here will serve as a new paradigm for developing RET systems.

## Acknowledgment

We are thankful to the following colleagues for providing plasmids: Xiaolan Yao (University of Missouri, Kansas City, USA) for plasmid pHis6-GB1 harbouring CERT PH domain; Gyun Min Lee (Korea Advanced Institute of Science and Technology, Daejeon, South Korea) for plasmid AAVS1-LP, and plasmid AAVS1-sgRNA-Cas9; Andras Balla and Laszlo.Hunyady (Semmelweis University, Budapest, Hungary) for plasmid RasBRET-tK. This work was financially supported by the USA National Niemann-Pick Disease Foundation and Deutsche Forschungsgemeinschaft, given to G.N. and C.A., respectively.

## Author contributions

C.A. initiated the project. C.A. and G.N. conceived the project. G.N. developed the concept, designed, and performed the experiments, and analyzed the data. C.A. and G.N. wrote the paper. Both authors take full responsibility for the content of the paper.

## Data availability

The relevant data are available from the corresponding authors upon request. The source data underlying Figs. 2c, 2e, 2g, 3a-d, and Figs. 1, 2a-b, 3a-b are provided as files.

## Declaration of competing interests

The authors declare no competing interests.

## Notes

### Competing Interest Statement

The authors have declared no competing interest.

