## Supplementary Information for "BRED: bioluminescence energy transfer to dye for monitoring ceramide trafficking in cell"

### Supplementary results

#### Intensities of the 535nm/480nm ratio of BRED donor and acceptor in HEK293 cells

The RET ratio of HEK293 cells was evaluated in presence of the donor substrate or acceptor. The results showed that adding 5 μM of CTZ-h (Fig. S1, dark yellow line) or 5 μM of BODIPY FL-Cer (Fig. S1, black line) to the HEK293 cells do not affect the signal emission at 480 nm and 520 nm. 520nm/480nm ratio values of this sample were similar to the level of noise signal of BRED 0.1 cells (Fig. 3, grey line) and HEK293 cells (Fig. 3, pink line).

#### Microscopy analysis of BRED output

Kim and Grailhe (2016) successfully employed a conventional widefield microscope for bioluminescence imaging of Nanoluc (max at 460 nm, 20 nm blue-shifted with respect to RLuc) [1]. The luminescence emitted by Nanoluc [2] is not a well-suited donor for BRED system. However, they could not observe a significant luminescence signal for cells transfected with plasmid encoding RLuc in combination with CTZ substrate (10 times less bright than CLZ-h). To monitor the bioluminescence signal generated using the energy donor of the BRED system, the bioluminescence imaging was performed for different cell lines using low BP filters 435-485 nm of a wieldfield bioluminescence microscope (see Supplementary Methods). As seen in Fig S2, no detectable signal was detected for control BRED cells, HEK293 cells treated 30 μM of CTZ-h (high concentration of CTZ-h was used to improve the bioluminescent signal level). HEK293 cells transfected with pGNPH in the presence of 30 μM of CTZ-h failed to display a bioluminescence signal using our widefield microscope. In contrast, a bioluminescence signal was detected in the case of BRED 0.1 exposed to 30 μM of CTZ-h. We then evaluated whether the bioluminescence emitted from BRED 0.1 in living cells could be employed to excite BODIPY FL-Cer. The bioluminescence images of the hRLuc (donor channel: BP 435-485 nm) and BODIPY FL-Cer (acceptor channel; BP 525-565) bioluminescence signals were performed (Fig. S2). BRED 0.1 cells exposed to 30 μM of CTZ-h showed a stronger signal in the donor channel, while BRED 0.1 cells exposed to 30 μM of CTZ-h and acceptor demonstrated a slightly stronger signal in the acceptor channel. BRED 0.1 exposed to CTZ-h, and BODIPY FL-Cer resulted in a signal-to-background ratio of ~7.2 (Fig S2 and see Supplementary Methods).

#### Flow cytometry method to assess the BRED output

The optical configuration for a combination of blue and YG channels of DB FACSMelody was setup. BRED 0.1 cells treated with BODIPY FL-Cer loaded into the system. The 527/32 nm filter is placed in a channel for yellow-green laser. The fluorescence values were obtained in blue (Fig.

S3a, left panel) and YG (Fig. S3a, right panel) channels. To improve the FACS generated signal through increasing the exposure time, 1 × $10^5$ cells were subject to the flow cytometry measurement for 10,000 events. Also, hRLuc-CTZ-h donor has a luminescence half-life of ~ 22 min. CTZ-h was added to the cells at a later stage, just prior to their passage through the laser beam, to improve the strength of signal output. After all optimizations, the fluorescent value for BRED 0.1 cells treated with BODIPY FL-Cer and CTZ-h were measured in the blue (Fig. S3b, left) and YG (Fig. S3b, right). Here, a clear shift to the right (Fig. S3, right panel) was observed for the BRED 0.1 cells, when the substrate was added. In contrast, there was no difference for the fluorescent output in the blue channel (Fig. S3, left panel). It is therefore evident that luminescence measurements can be performed using the DB FACSMelody flow cytometer.

### 1. Supplementary figures

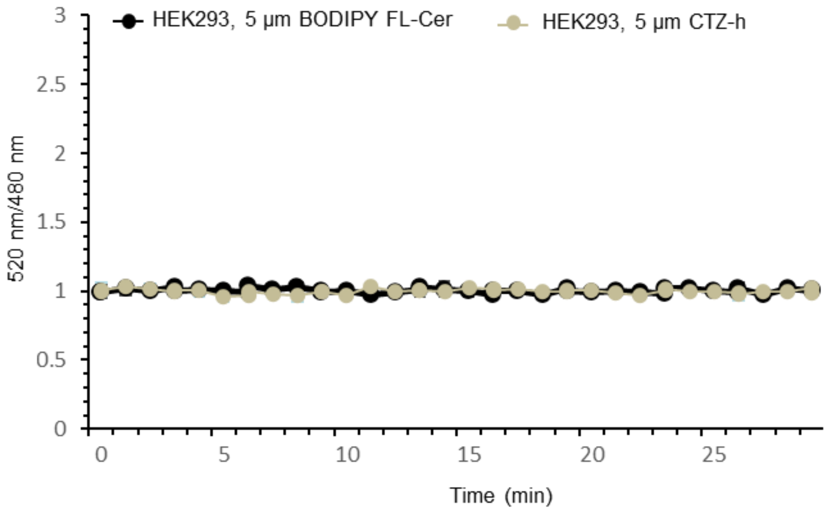


**Figure S1.** Intensities of the 535nm/480nm ratio of BRED donor and acceptor over the time in HEK293 cells.

Average 535nm/480nm ratio intensity measured every 45 s for 30 min for HEK293 cells treated with 5 μM of CTZ-h (dark yellow line) or 5 μM of BODIPY FL-Cer treatment (black line). Data are means ± SD from three biological replicates. BODIPY FL-Cer, borondipyrromethene fluorescein-ceramide; CTZ-h, Coelenterazine-h. Full data and source data are given in Data S4.

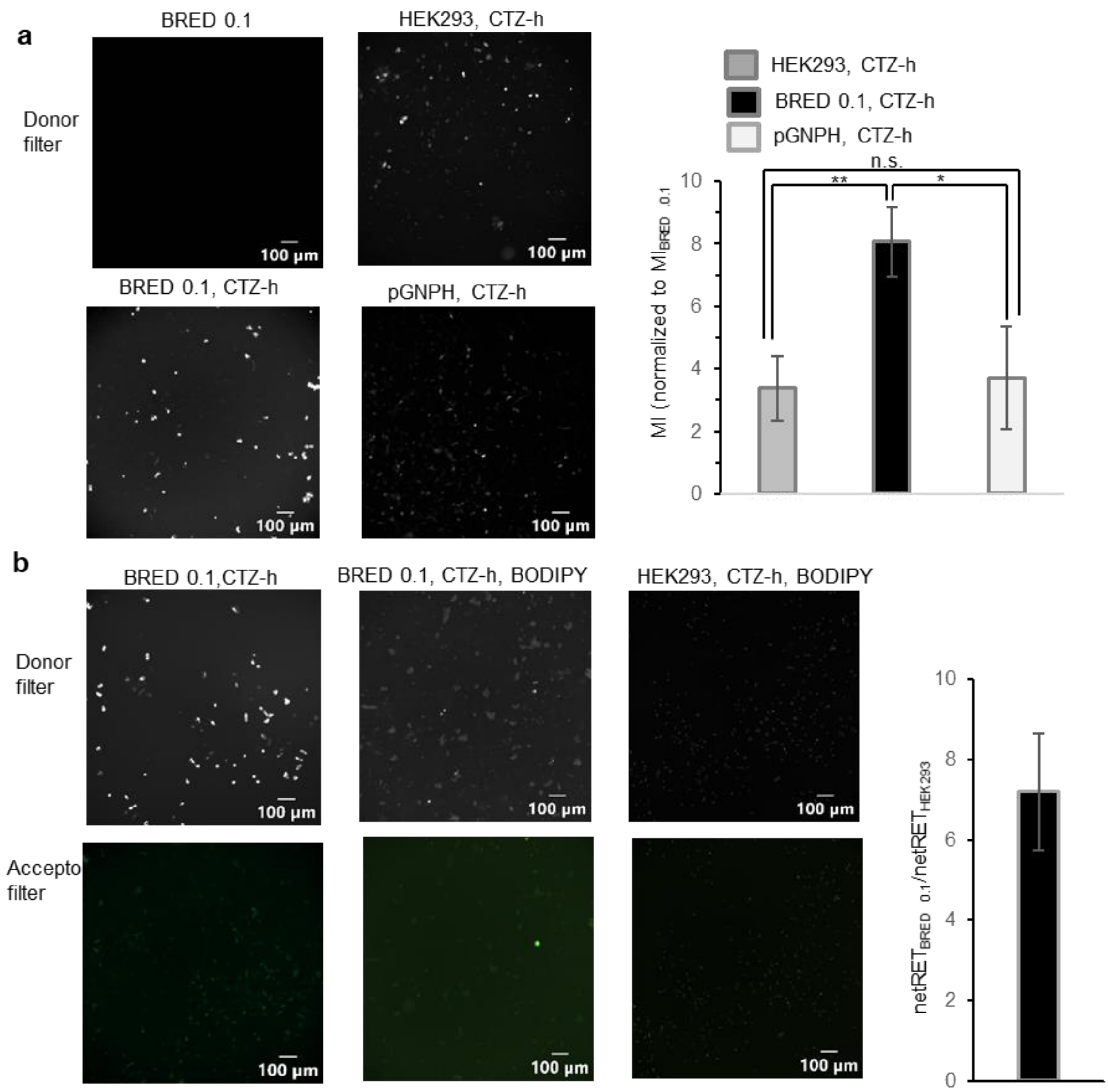


**Figure S2.** Microscopy analysis of BRED optical output.

a, Microscopy analysis of BRED donor. The bioluminescence images were acquired using a low banding pass filter 435-485 nm (Donor filter) for BRED 0.1 cells without CTZ-h substrate (top, left), HEK293 cells treated with 30 μM of CTZ-h (top, right), BRED 0.1 (bottom left) cells treated with 30 μM of CTZ-h, and HEK293 cells transfected with pGNPH and treated with 30 μM of CTZ-h (bottom, right). The graph shows the average optical intensity of three biological replicates obtained for HEK293 cells treated with CTZ-h (dark gray), BRED 0.1 cells treated with CTZ-h (black), and HEK293 cells transfected with pGNPH and treated with CTZ-h ((light gray). The BRED 0.1 cells showed higher optical output compared to HEK293 after adding CTZ-h. The optical output of HEK293 cells transfected with pGNPH was not significantly higher than HEK293 cells in presence of CTZ-h. The data were normalized to MI of BRED 0.1 cells without any treatment. b, Microscopy analysis of BRED donor and acceptor pair. The bioluminescence images were acquired using a low banding pass filter 435-485 nm (Donor filter), and high banding pass filter BP 525-565 (acceptor channel) for BRED 0.1 cells exposed to CTZ-h (left), BRED 0.1 cells exposed to CTZ-h and acceptor (middle) and HEK293 cells exposed to CTZ-h and acceptor (right). The graph shows the netRET[3] ratio between BRED 0.1 and HEK293 cells, both treated with CTZ-h and BODIPY FL-Cer. The experiment was performed in three biological replicates. CTZ-h, Coelenterazine-h; MI, mean intensity; Full data for Fig. S1a and b are given in Data S5a and b, respectively.

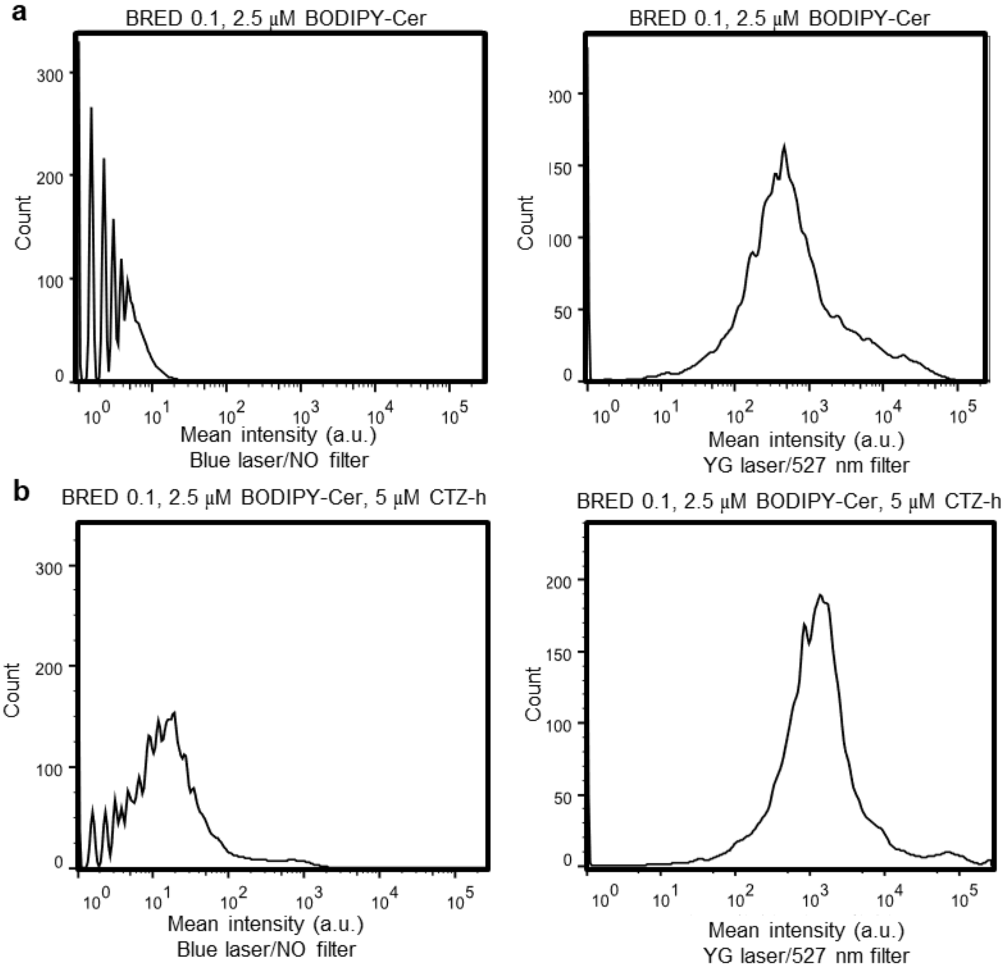


**Figure S3.** Flow cytometry histogram for optical configuration strategy.

After creating an optical configuration for the combination of blue and YG channels, MFI was measured for BRED 0.1 cells treated with 2.5 μM of BODIPY FL-Cer in blue (**a**, left panel) and YG (b, right panel), BRED 0.1 treated with 2.5 μM of BODIPY FL-Cer (**b**, left panel) and 5 μM CTZ-h (**b**, right panel). FACS data are representative of the mean ± SD of the fluorescence intensity obtained from three independent cultures and determined in three technical independent experiments. a.u., arbitrary unit, MFI, mean fluorescent intensity; YG, Yellow-green. Full data for Fig. S3a and b are given in Data S6a and b, respectively.

### Supplementary methods

Confocal microscope for BRED assay

To mission spectral scans of the BRED 0.1 and HEK293 cells expressing recombinant hRLuc, the cells were prepared in 8 well μ-slides as described in **Material and methods**. The experiment was performed using a widefield fluorescence IX83 microscope from Olympus, equipped with Orca Flash 4.0 V2 camera and objective UPLSAPO 10x lens with a 0.4 numerical aperture (NA). To remove a light source for widefield fluorescence microscopy required for focusing purposes, we first fixed the local position for slides. All options of cool light-emitting diode (LED) PE4000 were changed to “off”. The bioluminescence images of HEK293 non-transfected cells (background) were acquired and did not observe considerable light pollution from the equipment that could be detected on BP filters 410–480 nm and 525-565 nm. The bioluminescence images of hRLuc donor and the BODIPY FL acceptor emission for samples were acquired sequentially using low and high bandpass (BP) filters 435-485 nm; 525-565 at a confocal fixed position. The optical signals obtained from the donor, and acceptor filters for the HEK293 and BRED 0.1 cells exposed to CTZ-h and BODIPY FL-Cer were used to calculate the net BRET[1]. The bioluminescence signals from cells expressing the donor molecule only (BRED 0.1) using donor and acceptor filters were used to calculate the correction factor (cf) [3]. Using the cf, we corrected the bleed-through signal emitted from the donor hRLuc to the bioluminescence acceptor. Signal-to-background ratio for BRED 0.1 exposed to CTZ-h, and BODIPY FL-Cer was calculated using its bioluminescence divided by the average signal of images obtained with HEK293 cells in presence of CTZ-h, and BODIPY-FL. The bioluminescence quantification was performed using OlyVIA software.
